# Isolation by distance promotes strain diversification in the wild mouse gut microbiota

**DOI:** 10.1101/2025.09.15.676373

**Authors:** Brian A. Dillard, Jon G. Sanders, Azhar P. Husain, Kelsey M. Yule, Andrew H. Moeller

## Abstract

Bacterial species within the mammalian gut microbiota exhibit considerable strain diversity associated with both geography and host genetic ancestry. However, because geography and host ancestry are typically confounded, disentangling their contributions to the diversification of gut bacterial strains has remained challenging. Here, we show through joint profiling of gut bacterial and mitochondrial genomes from wild-living populations of deer mice (*Peromyscus maniculatus*) sampled across the United States that isolation by distance (IBD) drives gut bacterial strain diversification independently of the effects of host ancestry. Analyses revealed significant IBD in 27 predominant gut bacterial species, including members of the Muribaculaceae and Lachnospiraceae, but limited evidence for co-inheritance of gut bacterial genomes with mitochondria during the diversification of extant mouse populations. Gut bacterial species capable of forming spores exhibited reduced IBD independently of phylogenetic history, indicating that adaptations facilitating bacterial dispersal can mitigate the geographic structuring of strain diversity. These results show that the diversification of gut bacterial strains within rodent species has been mediated by geographic separation of host populations rather than host genealogical divergence.

## Main

Strain-level bacterial diversity in the mammalian gastrointestinal tract varies systematically among host individuals, populations, and species^1–5^. Despite the importance of strain-level variation in the gut microbiota for bacterial and host phenotypes^6–9^, the processes that drive strain diversification are not clear. Previous work in humans and other mammals has shown that strain divergence between hosts is associated with both host genetics and geographic distance^2,3,10–12^. These associations are consistent with the diversification of strains alongside host genealogical lineages, i.e., isolation by host (IBH), such as that mediated by maternal transmission of gut bacteria^13–15^. However, the associations are also consistent with isolation by distance (IBD), in which limits on bacterial dispersal imposed by geographic distance reduce gene flow between separated populations^2,16^. In most prior strain-resolved studies of the mammalian microbiota, the genetic relatedness of the host populations surveyed has mirrored their geographic distributions^3,12,16^, precluding quantification of the independent contributions of IBH and IBD to gut bacterial strain diversification.

A recent study of African apes, whose evolutionary relationships and geographic distributions have been uncoupled by migration throughout equatorial Africa, showed that gut bacterial strain divergence among these host species reflects host-species relatedness rather than geography^17^. However, the independent influences of IBH and IBD on the diversification of strains over more recent timescales, such as those that separate genealogical lineages or populations within host species, have not been quantified. For example, studies of humans have shown that gut bacterial strains have diversified in parallel with host populations^12^, a pattern consistent with IBH, but because human ancestry reflects geographically structured dispersal out of Africa, this pattern could have arisen from IBD in the absence of IBH^16,18^.

Here, we leveraged widespread geographic sampling of wild-living deer mice (*Peromyscus maniculatus*) to quantify the effects of IBH and IBD on the diversification of gut bacterial strains. Deer mice are widely distributed throughout North America, and prior population-genetic studies of this species have indicated gene flow between geographically separated populations^19,20^. Therefore, the biogeographic distribution of deer mice may enable quantification of the independent effects of IBH and IBD on gut bacterial strain diversification within a mammalian host species.

## Results

### Isolation by distance of deer mouse mitochondrial lineages

Before assessing the effects of IBD and IBH on bacterial strain diversification, we first asked whether deer mice exhibited evidence of IBD. We focused on maternal lineages, which, in mammals, have been proposed to mediate IBH of gut bacteria via maternal microbiota transmission^13^. We performed deep metagenomic shotgun sequencing on DNA extracted from 50 fecal samples collected from 43 deer mouse individuals at 13 field sites throughout the United States (Fig. 1A). To enable comparisons between host species, we also sequenced 44 fecal samples collected from 39 individuals representing three additional rodent species, *P. leucopus*, *Reithrodontomys megalotis*, and *Myodes gapperi*. Mapping reads against host reference genomes revealed that 124 million paired-end reads were derived from the rodent hosts with an average of 1.3 million paired-end reads per sample (SE = 174,000) (Table S1). Mapping these reads from deer mouse samples to the 16,322 bp deer mouse mitochondrial (MT) genome yielded an average breadth of coverage of 80.2% and an average depth of coverage of 15.4 reads per site.

**Figure 1.**
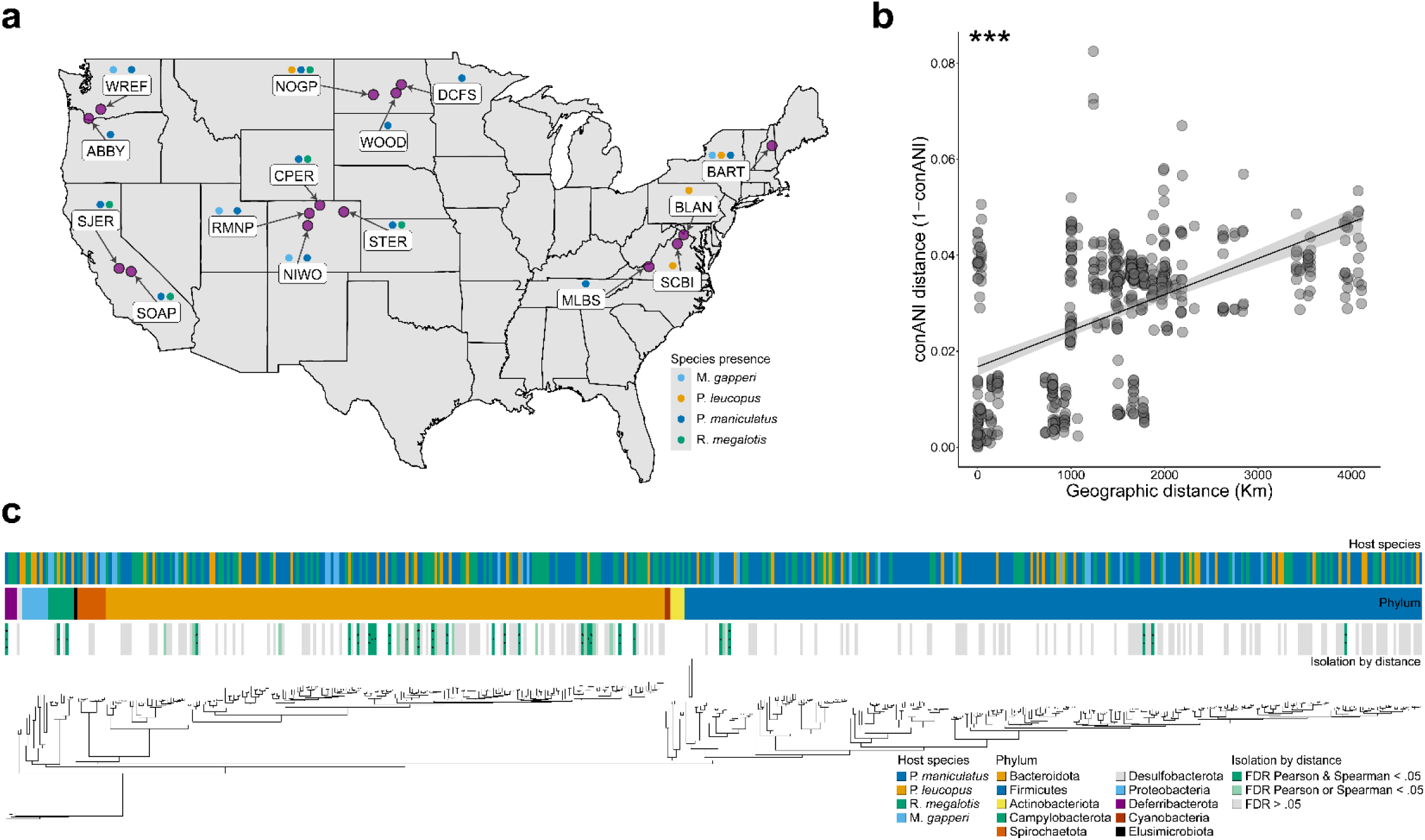
Isolation by distance shapes genetic diversity in deer mice and their gut microbiota. A) Map of the United States denotes the locations of field sites from which rodent fecal samples were collected. Field sites are labeled with site codes. Above each site label, colored points represent presence of rodent host species, as indicated by the key. B) Scatterplot shows isolation by distance of deer mouse maternal lineages. MT genetic distances and geographic distances are shown on the y- and x-axes, respectively. Line indicates best-fit trendline, and shading around the trendline shows standard error. Asterisk indicates p-value of Mantel test; *** < 0.001. C) Phylogeny of gut bacterial genomes (SGB representatives) assembled from the gut microbiota of deer mice and sympatric rodent populations. Branch lengths represent amino acid substitutions per site. Vertical bars marking tips indicate host species of origin (upper bar), phylum (middle bar), and significance testing for independent effects of IBH and IBD (bottom bar). In the bottom bar, grey rectangles mark SGBs passing filtering criteria and tested for independent effects of IBH and IBD, light green rectangles mark SGBs displaying significant evidence of IBD after accounting for IBH using either Pearson or Spearman MRM methods after false discovery rate (FDR) correction, and green rectangles mark SGBs displaying significant evidence of IBD after accounting for effects of IBH using both Pearson and Spearman MRM methods and after FDR correction. No SGBs displayed significant evidence of IBH after accounting for effects of IBD using both Pearson and Spearman MRM methods and after FDR correction. Asterisks in bars indicate MRM FDR-corrected p-values; * < 0.05, ** < 0.01, *** < 0.001.

Next, we used inStrain^5^ to calculate pairwise consensus average nucleotide identity (conANI) between MT genomes of all individuals. Using these data, we conducted a Mantel test to assess the strength of IBD on deer mouse MT genetic divergence. Deer mouse MT genetic distances (1 – conANI) were positively associated with geographic distances (Mantel’s r = .522, p value = .00001) (Fig. 1B), indicating significant IBD in deer mice. MT conANI between deer mouse individuals was, in every case, within the ANI ranges previously described for this species^21,22^, validating host species assignments of fecal samples. Significant IBD was also observed when deer mice from each individual field site were compared to mice from every other field site (Fig. S1), indicating that the significant effects of IBD were not exclusive to any one field site or subset of field sites. However, we observed considerable variation around the trendlines describing the relationships between host mitochondrial divergence and geographic distances (Fig. 1B; Fig. S1), enabling quantification of independent effects of IBD and IBH on gut bacterial strain diversification.

### Geography structures gut bacterial strain diversity in deer mice independently of maternal relatedness

To investigate the relative contributions of IBD and IBH to gut bacterial strain diversification in deer mice, we generated a set of reference gut bacterial metagenome-assembled genomes (MAGs) from these hosts. Sequencing of fecal samples from deer mice and other host species yielded a total of 2.68 billion bacterial paired-end reads, with an average of 28.5 million paired-end reads per sample (SE = 913,000) (Table S1). Using the filtered bacterial reads from each host, we used a reference-free metagenome assembly and genome binning approach^23–26^ to produce 1,012 *de novo* chimera-free MAGs that met draft quality thresholds of at least 50% completeness and less than 5% contamination^27,28^, including 538 MAGs from deer mice (Table S2). Analyses of the 1,012 MAGs identified 492 95% average nucleotide identity (ANI) clusters, i.e., species-level genome bins (SGBs). We used bowtie2^29^ to map quality-filtered bacterial reads from each fecal sample back to the reference set of SGBs. Then, for every sample, we used inStrain^5^ to profile the allelic diversity found within each of the SGBs. On average, 50% (SE = 1.3%) of bacterial reads per deer mouse sample mapped to the SGB reference genomes, indicating that the SGB reference genomes were broadly representative of the bacterial diversity in the metagenomes.

Next, we quantified the independent effects of IBD and IBH on strain divergence between host individuals. For each of the gut bacterial species detected in deer mice, we tested for associations of inter-host strain divergence with geographic distance while controlling for host MT distance and with MT distance while controlling for geographic distance. Of the 492 SGBs, 197 were present in at least three deer mouse individuals and in at least two populations. These 197 SGBs had a mean depth of coverage of 6.7 (SE = 0.57), and a mean breadth of coverage of 60% (SE = 1%), providing sufficient information for genome-wide comparisons of strains between individual hosts (Table S2). A maximum likelihood phylogeny of the 197 SGBs was inferred using GTDB-Tk^30^ and the Bac120 single copy core gene alignment is presented in Fig. 1C. This phylogeny recovered reciprocal monophyly between 10 gut bacterial phyla, including those known to be most abundant in the mammalian gut microbiota (e.g., Firmicutes and Bacteroidota). For each of these 197 SGBs, we calculated strain genetic distance (1 – conANI as measured by inStrain) between every pair of samples. We then used multiple regression on distance matrices (MRM) with both Spearman and Pearson correlation methods to test for independent effects of geographic distances and host MT genetic distances on strain genetic distances within each SGB.

After Benjamini-Hochberg correction for multiple comparisons across SGBs, 27 SGBs displayed strain diversification significantly structured by IBD independently of IBH based on both Pearson and Spearman correlation coefficients (MRM, FDR-corrected p-value < 0.05) (Fig. 1C; Fig. 2; Table S3). In contrast, no SGBs displayed significant evidence of IBH independent of IBD in both analyses (MRM, FDR-corrected p-value > 0.05 in every case) (Table S3). To visualize these results, we used generalized least squares (GLS) to model the relationship between strain genetic distance (1 – conANI) and geographic or host MT distance, then plotted the residuals derived from these GLS analyses against geographic distance (Fig. 2) or host MT distance (Fig. S2).

**Figure 2.**
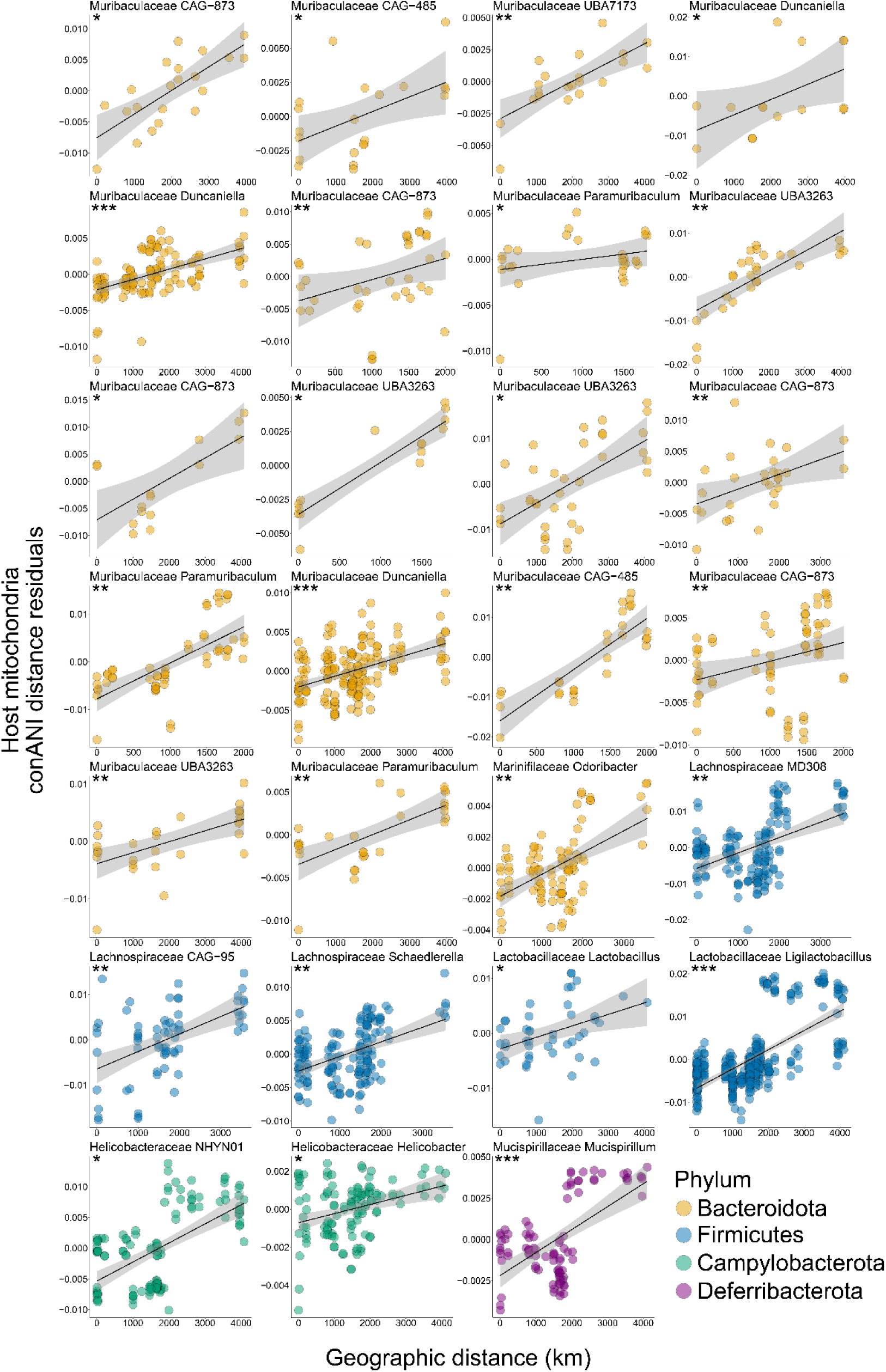
Isolation by distance drives strain diversification within gut bacterial species. Scatterplots and trendlines show significant positive relationships between SGB conANI distances (residualized against host MT distances) and geographic distances for individual SGBs derived from *P. maniculatus* hosts. SGB ANI distances were residualized against residuals from a GLS model with the formula: SGB ANI distance ∼ host mitochondrial ANI distance. Each facet represents an SGB, with colors denoting the phylum classifications for each SGB, as indicated by the key. Shading around each trendline shows standard error from best-fit linear model. Asterisks indicate MRM Pearson FDR-corrected p-values; * < 0.05, ** < 0.01, *** < 0.001.

SGBs structured by IBD independently of IBH represented predominant taxa in the deer mouse gut microbiota, including Muribaculaceae, Lactobacillaceae, Lachnospiraceae, and Helicobacteraceae. Mean depth of coverage did not significantly differ between the 27 SGBs affected by IBD and those not affected by IBD, suggesting that the relative abundance of SGBs did not bias the detection of IBD (permutation t-test p-value > 0.05). Combining all SGBs into a single analysis also confirmed that strain genetic distances were significantly and strongly positively associated with geographic distances independently of host genetic distances, indicative of IBD (Fig. S3A). In contrast, host genetic distances had relatively modest effects on strain genetic distances after controlling for the effects of geography (Fig. S3B). Cumulatively, these results indicate that IBD, rather than IBH, has driven gut bacterial strain diversification in deer mice.

### Sporulation ability mitigates the effects of IBD on gut bacterial strain diversification

Given that some SGBs displayed significant evidence of IBD whereas others did not, we next tested whether bacterial traits explained variation in the strength of IBD across SGBs. We hypothesized that sporulation ability and motility may lessen the contribution of IBD to strain diversification by increasing bacterial dispersal ability, based on previous studies that have identified associations between dispersal-related traits and rates of gut bacterial transmission^15,31^. We used gene ontology annotations^32,33^ to determine the abundances of sporulation and motility genes in each of the 197 deer mouse SGBs. We then used phylogenetic independent contrasts to assess the effects of sporulation or motility gene abundances on the strength of IBD, measured as the MRM correlation coefficient between strain genetic distances (1 – conANI) and geographic distances after controlling for host MT distances. Results revealed that, after controlling for bacterial evolutionary history using phylogenetic independent contrasts (PICs), sporulation gene abundances were significantly and negatively associated with the strength of IBD (permutation linear model p-value for negative slope = 0.032) (Fig. 3A). In contrast, motility gene abundances were not associated with the strength of IBD. In addition, we used Traitar^34^ to test whether a broad suite of bacterial traits were associated with the strength of IBD. These analyses, which analyzed 54 well-annotated bacterial traits, confirmed the negative association between the presence or absence of sporulation ability and the strength of IBD (GLS, p-value < 0.05) (Fig. 3B; Fig. S4; Table S4). Together, these findings indicate that sporulation ability mitigates the effects of IBD on strain diversification within the rodent gut microbiota.

**Figure 3.**
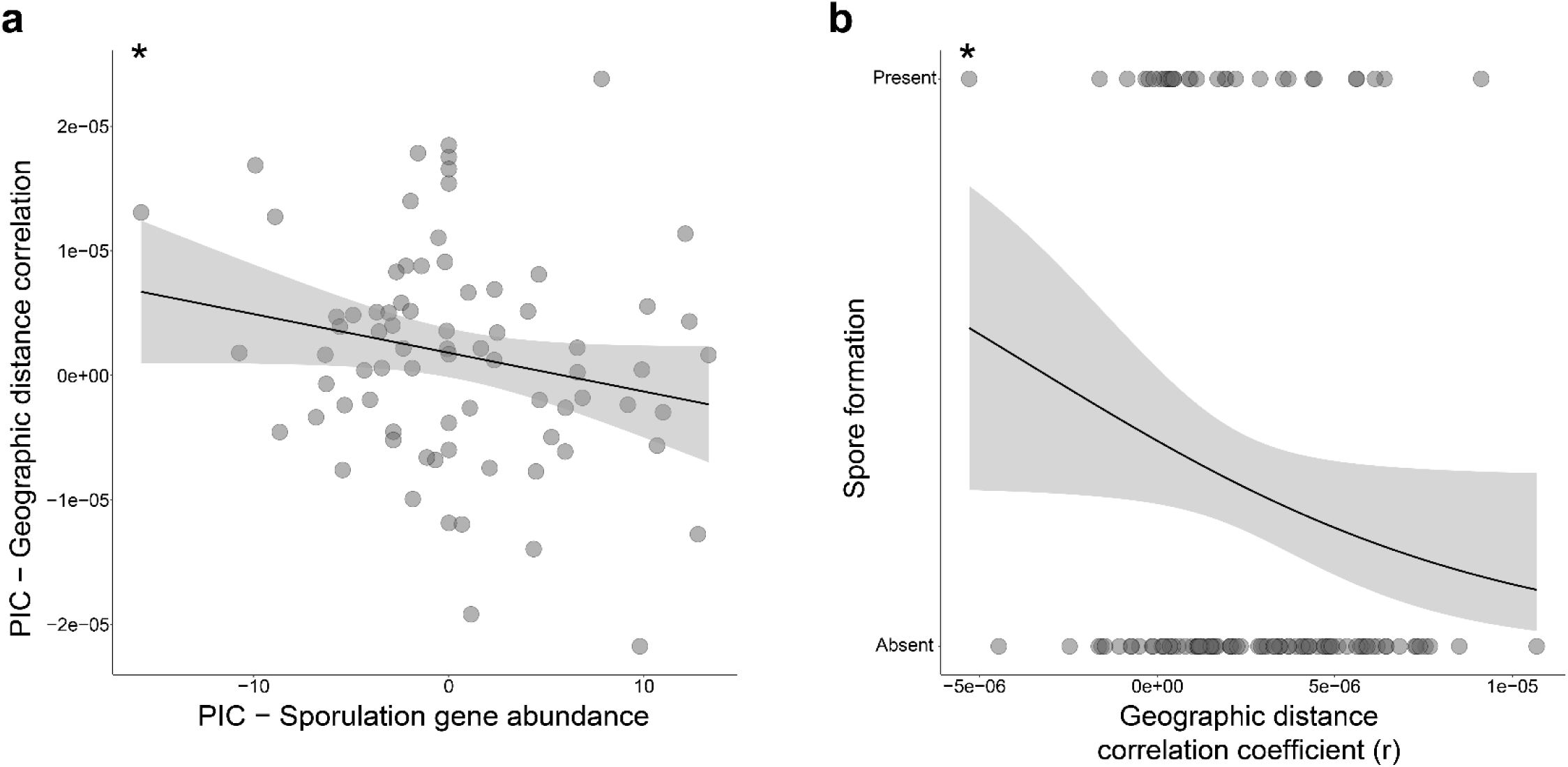
Sporulation ability mitigates effects of isolation by distance on strain diversification. A) Scatterplot shows negative relationship between the abundance of spore forming genes in the genomes of gut bacterial species (x-axis) and the strength of IBD (y-axis), after accounting for bacterial phylogenetic history using phylogenetically independent contrasts (PICs). The strength of IBD (y-axis) was measured by the geographic distance correlation coefficient estimated by MRM after accounting for host MT distances. Trendline shows best-fit linear regression. Shading around trendline denotes standard error. Asterisks indicate permutation linear model p-value for negative slope; * < 0.05. B) Logistic regression plot shows negative relationship between the strength of IBD for SGBs (x-axis) and the presence or absence of sporulation ability inferred by Traitar (y-axis). Asterisks indicate p-value of GLS model; * < 0.05.

### Maternal relatedness structures strain diversity between, but not within, rodent genera

The results that IBD, not IBH, explained the biogeographic distribution of gut bacterial strain diversity within host species (Figs. 1 and 2) contrast the results of studies that have found evidence for strain IBH between mammalian species after controlling for the effects of geography^17^. Moreover, prior work in primates and rodents has shown that host species living in the same geographic locale (i.e., in sympatry) maintain compositionally distinct gut microbiota that reflect their hosts’ phylogenetic relationships^35,36^. This disparity suggests that the effects of IBH on the generation and maintenance of strain diversity in the gut microbiota may only manifest over timescales that separate host species. To test this idea, we analyzed 29 SGBs from deer mice that were also detected in sympatric heterospecific host populations from locations where deer mice co-occurred with either *P. leucopus* or *R. megalotis*. *M. gapperi* was excluded from this analysis due to a lack of shared SGBs with deer mice, which could be explained by less intensive sampling of *M. gapperi* and greater evolutionary divergence from deer mice, relative to *P. leucopus* or *R. megalotis*. These 29 SGBs had a mean breadth of coverage of 58% (SE = 2.5%) and a mean depth of coverage of 10.1 (SE = 2.58), providing sufficient information for genome-wide comparisons of strains within and between host species (Table S2).

For each SGB shared by sympatric hosts and present in at least two individuals of each host species, we compared mean microbial genetic distance (1 – conANI) across three host pairings: conspecific (*P. maniculatus* vs. *P. maniculatus*), congeneric (*P. maniculatus* vs. *P. leucopus*), and heterogeneric (*P. maniculatus* vs. *R. megalotis*). These comparisons revealed that gut bacterial strains within shared SGBs were more genetically divergent between sympatric heterogeneric hosts than between sympatric conspecific or congeneric hosts (permutation t-test of mean genetic distance per SGB, p-value < 0.05) (Fig. 4). These analyses identified 7 SGBs, including representatives from the families Anaerotignaceae, Muribaculaceae, Lactobacillaceae, and Lachnospiraceae, for which heterogeneric host pairs displayed significantly elevated strain genetic distances relative to congeneric pairs (permutation t-test, FDR p-value < 0.05). A table summarizing results of SGB comparisons between conspecific hosts and between host species pairs is present in Table S5. Cumulatively, these results show that, although IBH has negligible effects on the distribution of gut bacterial strain diversity within host populations and species, IBH maintains differences in strain diversity between sympatric host species.

**Figure 4.**
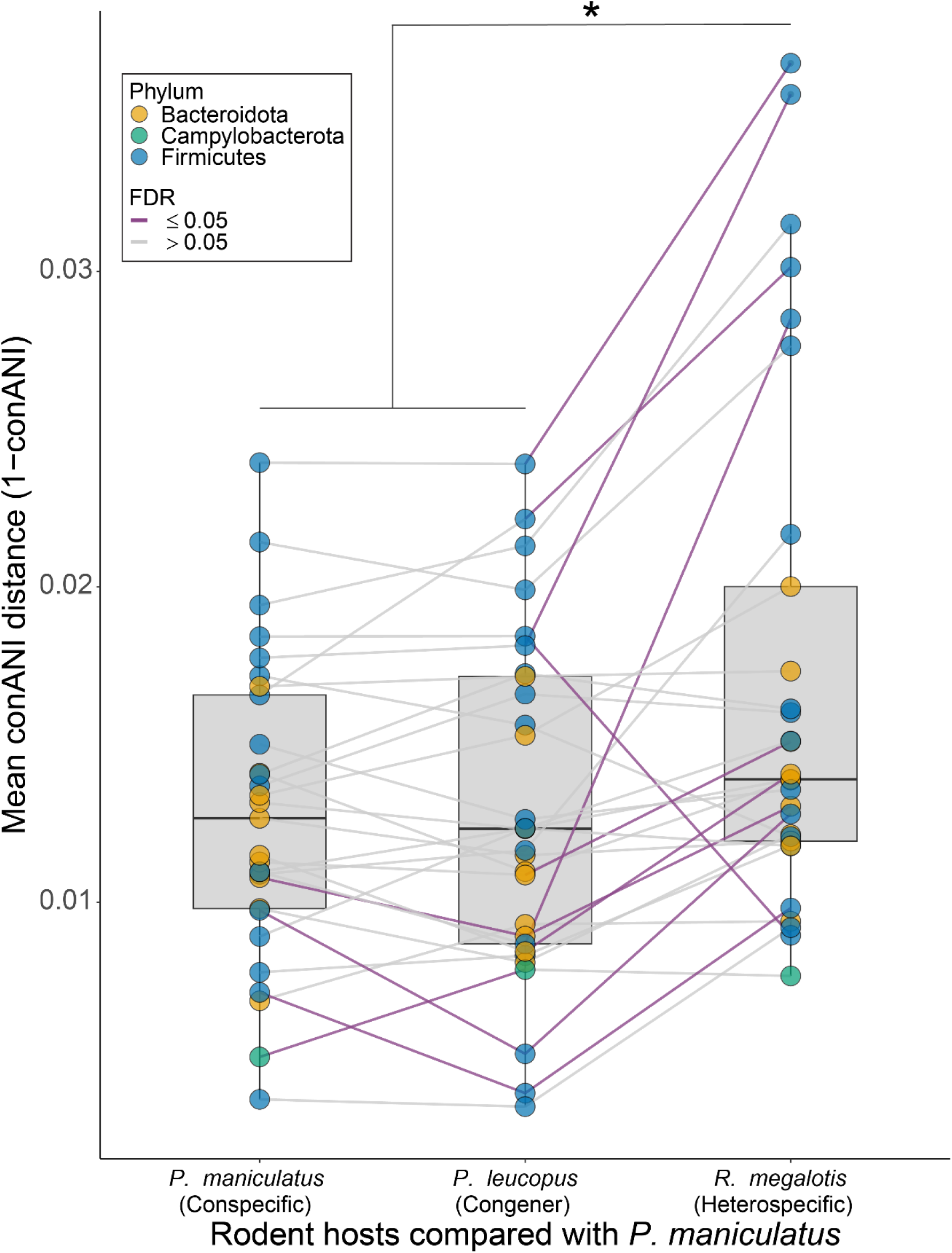
Sympatric host genera maintain genomically distinct gut bacterial strains. Boxplots show the genomic divergence of strains from shared SGBs between individual hosts derived from three different sympatric rodent host pairings: *P. maniculatus* vs. *P. maniculatus* (left), *P. maniculatus* vs. *P. leucopus* (center), and *P. maniculatus* vs. *R. megalotis* (right). Each point represents the mean of comparisons for an SGB with color denoting phylum classifications, as indicated in the key. Lines connect the same SGB across host pairings with purple lines indicating significant differences between pairings (permutation t-test, FDR < 0.05). Asterisk denotes significant difference between means of pairs of boxplots (i.e., significant difference between the mean of the means of SGB genetic distances between pairings) based on permutation t-test p-value; * < 0.05.

## Discussion

Comparative analyses of the allelic variation present in metagenomes and mitochondria of wild rodent populations from across the United States provided insights into the processes driving the diversification of gut bacterial strains within host species. Results indicated that gut bacterial strains have not been obligately inherited within or diversified alongside host maternal lineages; instead, the diversification of gut bacterial strains has been mediated primarily by dispersal limitation imposed by geographic distance. Most gut bacterial species whose strain diversity was structured by IBD belonged to the phylum Bacteroidota (e.g., Muribaculaceae spp.) (Fig. 1). In humans, variation within gut-associated Bacteroidota species has been previously associated with both host genetic ancestry and geography^12^, though prior work was not able to disentangle the independent contributions of these factors to strain diversification^3,12^. Our results indicate that geographic separation promotes the diversification of gut bacterial strains independently of the effects of host ancestry (Figs. 1 and 2; Fig S2).

Interestingly, gut bacterial species that lacked the ability to form spores were most strongly affected by IBD. Across bacterial species, sporulation ability was negatively associated with the strength of IBD, even after controlling for bacterial phylogenetic history. Previous work has shown that spore forming ability, which mediates bacterial dispersal, is associated with increased rates of transmission of gut bacterial strains between inbred lines of laboratory mice^37^. Our findings show that sporulation can mitigate the geographic structuring of strain diversification within wild living mammalian species, likely due to the positive effects of sporulation on microbiota dispersal rates.

We observed no significant effects of IBH on the biogeographic distributions of gut bacterial strains within deer mouse SGBs after accounting for geography. Past work in laboratory mice has shown that gut bacteria can be inherited through maternal lines over decadal timescales^15^, but our results from wild-living populations indicate that gut bacterial strains have not been obligately inherited through maternal lineages over the timescales of the diversification of extant deer mouse populations. In contrast, comparing strain diversity of the deer mouse microbiota with that of sympatric heterospecific and heterogeneric host populations indicated significant effects of IBH in maintaining strain divergence between rodent genera (Fig. 4). Together, these results establish a stepwise process through which host-microbe symbioses diversify in mammals: IBD drives gut bacterial strain diversification within host species, whereas effects of IBH emerge over longer evolutionary timescales that separate host genera.

## Methods

### Sample collection

Small rodent fecal samples from *Peromyscus maniculatus*, *Peromyscus leucopus*, *Reithrodontomys megalotis*, and *Myodes gapperi* were collected by the National Environmental Observatory Network (NEON)^38,39^ across 13 field sites distributed throughout the continental United States. Rodents were captured using Sherman traps arrayed in a 10 x 10 grid with 10m between each trap and baited with a sterilized seed mixture consisting of 35% sunflower seeds and 65% millet. For each captured rodent, age, reproductive status, sex, and standard measurements were recorded. Host species was determined in the field based on diagnostic morphological traits, then confirmed with MT genotyping from metagenomic data (discussed below). Fresh fecal samples were collected with either forceps or by scooping the sample into a cryovial directly. Samples were then stored at -80 °C until requested, after which they were shipped to Cornell University on dry ice and again stored at -80 °C until DNA extraction.

### DNA extractions, library preparation, and Illumina shotgun metagenomic sequencing

DNAs were extracted from the rodent fecal samples using the bead beating based Qiagen DNeasy Powerlyzer PowerSoil kit according to the manufacturer’s instructions. Extracted DNAs were submitted to the Cornell Biotechnology Resource Center for TruSeq equivalent DNA library preparation. Library preparations were then sent to the UC Davis DNA technology core for paired-end short read sequencing on an Illumina NovaSeq sequencer.

### Assembly of rodent MAGs

Rodent derived MAGs were assembled using a metagenomic snakemake v8.3.2^40^ pipeline available at https://github.com/CUMoellerLab/sn-mg-pipeline. Adapters were trimmed from Illumina paired-end sequencing reads using Cutadapt v1.13^41^ and then checked for quality with FastQC v0.11.9^42^. Using bowtie2 v2.4.5^29^, processed sequencing reads were mapped to the most closely related host reference genome available from NCBI (*P. maniculatus* & *R. megalotis*: GCA_003704035.3; *P. leucopus*: GCA_004664715.2; *M. gapperi*: GCA_902806735.1). Reads that mapped to their respective host genome were removed from all downstream bacterial analysis using SAMtools v1.16.1^43^, as they were likely derived from the host. Filtered reads were then assembled into contigs with Megahit v1.2.9^26^ and the resulting contigs were checked for quality using Quast v5.2.0^44^.

Prior to binning, we utilized a prototype selection^45^ methodology for cross sample mapping of samples derived from the same host species. Jaccard distance matrices were created for samples derived from each host species using SourMash v4.8.8^46^, and samples that were of sufficient quality and represented the most genetic diversity had their reads mapped to the contigs of all other samples derived from the same host species using Minimap2 v2.24^47^. Mapping files were combined into coverage tables and contigs were binned using CONCOCT v1.1.0^25^, MaxBin 2.0 v2.2.7^24^, and MetaBAT 2 v2.15^23^. DAS Tool v1.1.7^48^ was then used to choose the highest quality non-redundant genome bins from the three different binning algorithms. MAGs that had less than 50% completion, more than 5% contamination, or high levels is chimerism as detected using default parameters with CheckM2 v1.0.2^27^ and GUNC v1.0.6^28^ were removed from downstream analysis.

### Taxonomic classifications of MAGs and inference of a genome-resolved phylogenetic tree

For each MAG that passed the quality filters, the classify workflow from GTDB-TK^30^ was used for taxonomic classification. Core genes from the Bac120 collection were identified using ‘identify’ and then concatenated for each MAG. Concatenated core genes were aligned using the ‘align’ function, with the resulting alignment run through the ‘classify’ function which places each MAG onto the GTDB-TK reference tree. Additionally, the GTDB-TK core gene alignment was used to infer a maximum likelihood phylogenetic tree of the MAGs using IQ-tree^49^.

### Intraspecific analyses of bacterial diversity

Rodent MAGs were dereplicated at a 95% ANI cutoff with dRep v3.0^50^, producing a set of microbial species level genome bins (SGBs). The highest quality MAG from each SGB was selected as the representative reference genome. The complete set of SGB reference genome fasta files was concatenated. Processed reads from each sample were competitively mapped against this concatenated reference using bowtie2^29^. Microdiversity analyses were performed with InStrain v1.8.0^5^ using the profile command in --database_mode, which sets the minimum read ANI to 0.92 and the minimum genome coverage to 1.0. Pairwise comparisons between all SGB profiles were then conducted with InStrain compare. Reads mapping to their respective host mitochondrial genomes during the read processing steps were also profiled and compared with InStrain using the same ANI and coverage. For rodent hosts with multiple fecal samples, we retained only one sample per host based on the quality of the microdiversity analyses, with higher average number of compared bases and SNPs being the main criteria for inclusion.

### Isolation by distance of deer mouse maternal lineages

Genetic distances between conspecific rodent hosts was calculated as 1 – conANI using InStrain compare. ConANI compares the major allele between two samples at every shared site. Geographic distance was computed using the haversine formula, implemented in R with the distHaversine command in the geosphere package, which calculates the distance between two points on a sphere using latitude and longitude. Latitude and Longitude coordinates for each rodent sample were based on the NEON sampling locations. The correlation between genetic and geographic distances was calculated using a Mantel test with the Pearson method and 10,000 permutations as implemented in Vegan v2.6-6.1^51^.

### Host and geographic effects on strain diversity within SGBs derived from P. maniculatus

We assessed the independent effects of host mitochondrial genetic distance and geographic distance on strain genetic distance within SGBs derived from *P. maniculatus*. Analyses were restricted to the 197 SGBs detected in at least three individual hosts and at least two populations. For each SGB, we performed multiple regression on distance matrices (MRM) using both Spearman and Pearson methods as implemented in the *ecodist* package v2.1.3^52^. MRM was run with up to 10,000 permutations, depending on the number of samples in which each SGB was found, using the formula (strain genetic distance ∼ host mitochondria genetic distance + geographic distance). Resulting *p*-values were corrected for multiple testing using the Benjamini–Hochberg procedure.

To visualize these results, we calculated the correlation between strain genetic distance (1 – consensus ANI) and both host mitochondria genetic distance and geographic distance using GLS models with the formulas (strain genetic distance ∼ geographic distance) and (strain genetic distance ∼ host mitochondria genetic distance) respectively. We then extracted the residual effects from each model and plotted them against host MT or geographic distance respectively.

### Testing for association between dispersal-related traits and isolation by distance

SGBs with an estimated completeness of at least 90%, as assessed by CheckM2 v1.0.2^27^, and present in at least four samples were functionally annotated using Prokka with Prodigal^53^. A curated list of bacterial genes associated with bacterial processes relating to sporulation and motility were obtained from the gene ontology knowledgebase^32,33^ (categories: 0043934 and 0048870, respectively) and filtered to retain only those genes with UniProt identifiers present in at least one annotated SGB.

To account for shared evolutionary history, both sporulation gene abundances and SGB-specific Pearson correlation coefficients from prior MRM analyses were transformed using phylogenetic independent contrasts (PIC) via ape v5.8-1^54^. The phylogeny used for PIC calculations included all SGBs and was constructed using IQ-tree^49^, rooted at the midpoint, and pruned to match the set of SGBs included in this analysis. Following previous studies^29^, any sample with a PIC value identified as a statistical outlier, defined as any PIC 1.5 times the interquartile range above the third quartile or below the first quartile, was removed prior to linear modeling.

The relationship between the strength of each SGB’s association with geographic distance and either sporulation or motility gene abundances was assessed using a permutation based linear model implemented with permuco v1.1.3 using the formula (Gene abundance PIC ∼ Geographic distance correlation coefficient PIC -1) and 100,000 permutations.

### Testing for association between traits and isolation by distance with Traitar

For each SGB that was above 90% complete, bacterial traits were predicted using Traitar^34^, a machine learning phenotype classifier that uses phylogenetic histories to estimate the presence of 54 bacterial traits. Trait designations were only considered if both the phypat and the phypat + PGL classifiers were in consensus. We used GLS to measure the association between each of the bacterial traits and the Pearson MRM correlation coefficients.

### SGB genetic distances between sympatric hosts

To compare the extent of SGB sharing between *P. maniculatus* and other rodent hosts sampled at sympatric field sites, mean consensus ANI distance (1 – conANI) was calculated for each SGB found with at least two pairwise comparisons in each of three host comparisons: *P. maniculatus* vs. *P. maniculatus* (conspecific), *P. maniculatus* vs. *P. leucopus* (congeneric), and *P. maniculatus* vs. *R. megalotis* (heterospecific).

To evaluate whether genetic distances between *P. maniculatus* and other rodent hosts differed significantly from conspecific *P. maniculatus* comparisons, permutation t-tests were conducted using up to 10,000 permutations, depending on the number of samples. Two permutation tests were performed per SGB: one comparing conspecific *P. maniculatus* comparisons to congeneric *P. maniculatus* vs. *P. leucopus* comparisons, and one comparing congeneric *P. maniculatus* vs. *P. leucopus* comparisons to heterogeneric *P. maniculatus* vs. *R. megalotis* comparisons. The resulting p-values were adjusted using the Benjamini-Hochberg procedure.

## Data and Code Availability

All sequence data generated in this study has been deposited in the National Center for Biotechnology Information Sequence Read Archive under the BioProject accession PRJNA1306187. All bacterial metagenome-assembled genomes generated in this study are available at https://doi.org/10.5061/dryad.95x69p8z4. Code used to setup and run the SGB diversification analyses is available at https://github.com/briandill2/Strain_diversification.

## Acknowledgements

We thank Dr. Weiwei Yan for assistance with DNA extractions from rodent fecal samples. The National Ecological Observatory Network is a program sponsored by the U.S. National Science Foundation and operated under cooperative agreement by Battelle. This material uses specimens and/or samples collected as part of the NEON Program and provided by the NEON Biorepository at Arizona State University. Funding was provided by the National Institutes of Health grants R35 GM138284 and R01 DK139214 to AHM.

## Author Contributions

B.A.D designed the study, performed analyses, and wrote and edited the manuscript. J.G.S designed the study and edited the manuscript. A.P.H and K.M.Y facilitated sample selection and provisioning, and edited the manuscript. A.H.M supervised the project, designed the study, and wrote and edited the manuscript.

## Supplementary Materials

### Supplementary Figures

**Fig. S1.**
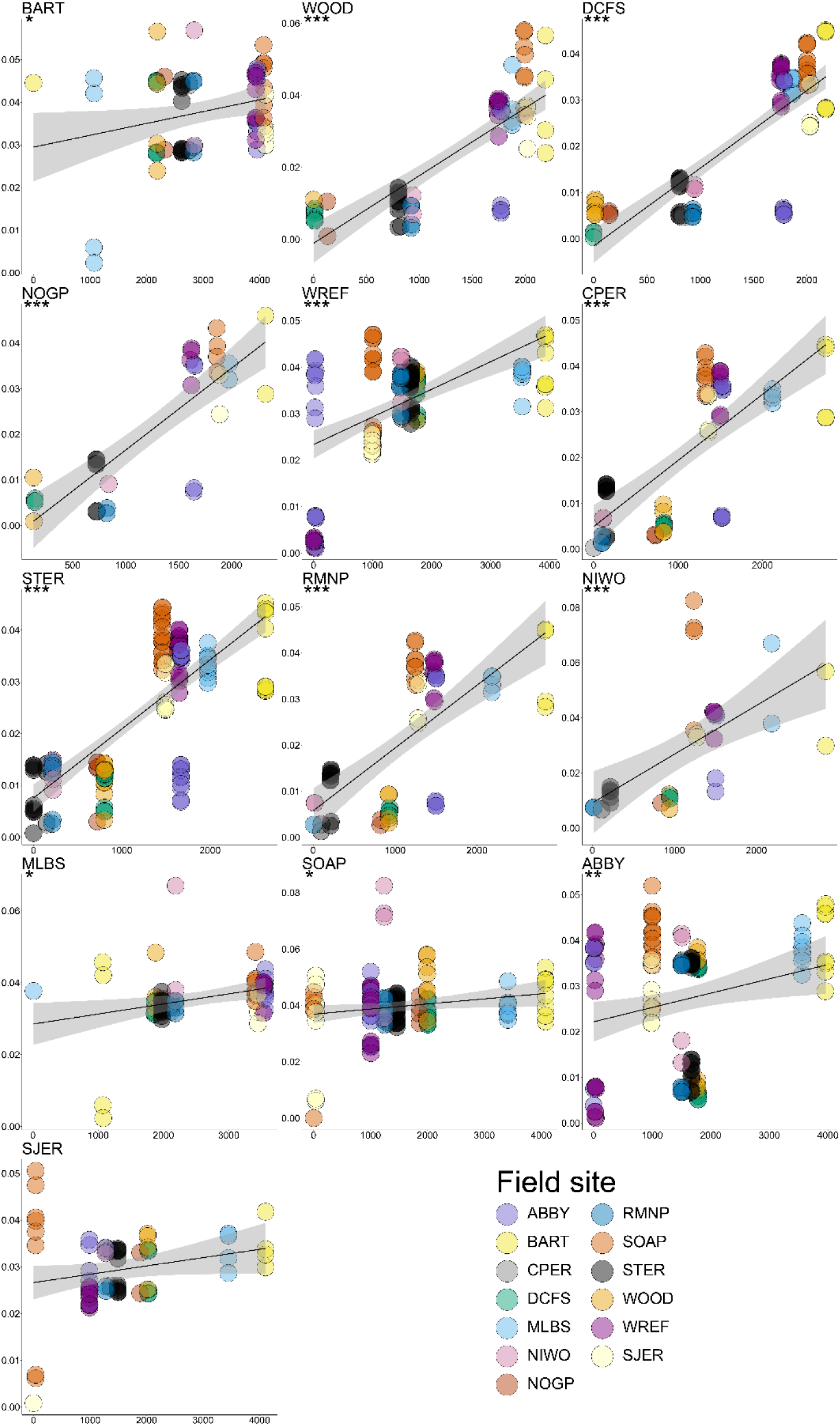
Consistent evidence for IBD in deer mice across field sites. Scatter plots show IBD of deer mouse MT lineages across field sites. MT genetic distances and geographic distances are shown on the y- and x-axes, respectively. Each facet corresponds to a focal field site for which comparisons including that field site are shown. Points represent pairs of deer mice, with color denoting the non-focal field site. Trendline and shading indicate best-fit linear regression with standard error.

**Fig. S2.**
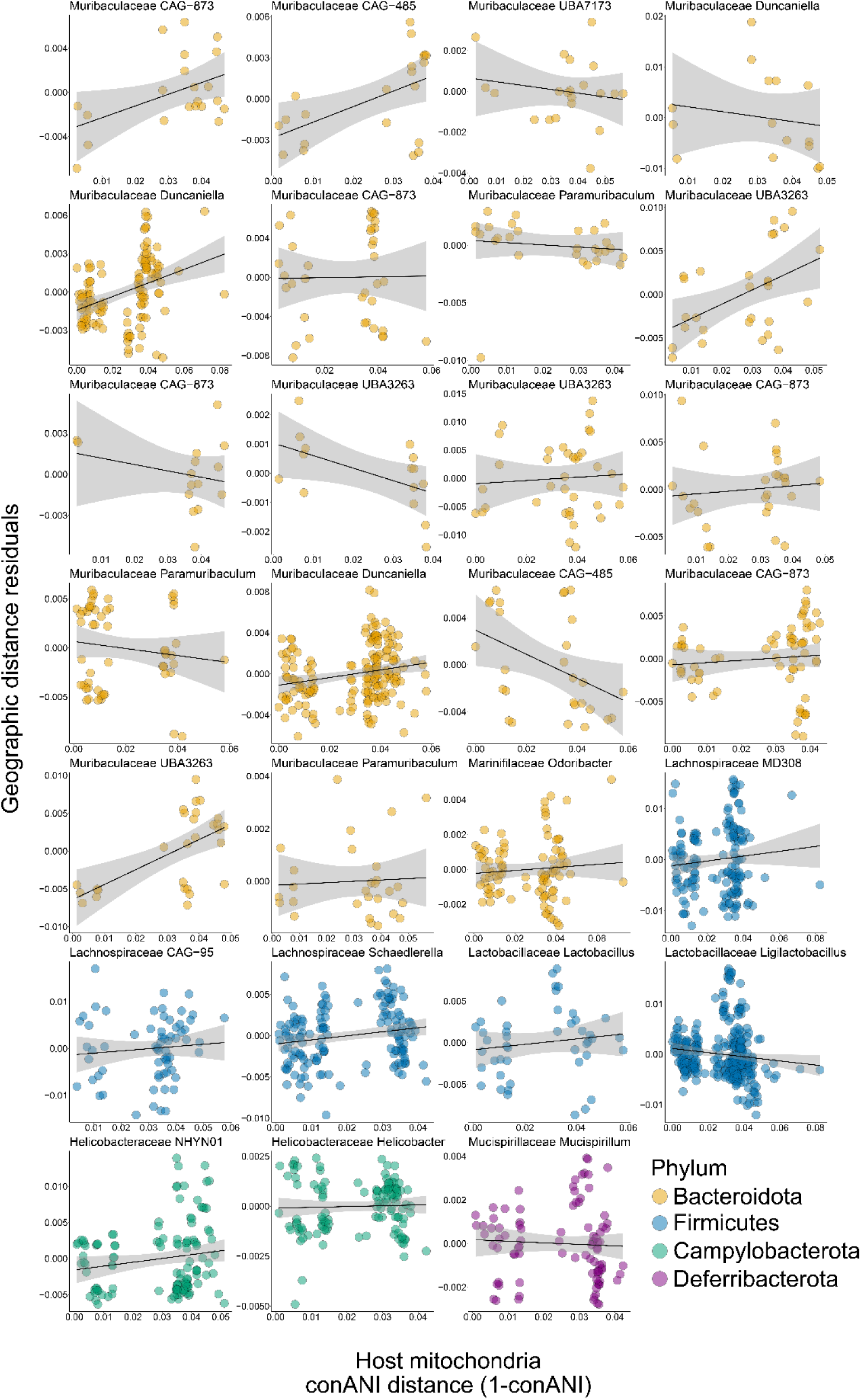
No effect of IBH on strain diversification within gut bacterial species. Scatterplots and trendlines show lack of relationships between SGB conANI distances (residualized against geographic distances) and host MT distances for individual SGBs derived from deer mouse hosts. SGB ANI distances were residualized against the residuals from a GLS model with the formula: SGB ANI distance ∼ geographic distance. Each facet represents an SGB. Colors denote the phylum to which each SGB belongs, as indicated by the key. Shading around each trendline shows standard error from the best-fit linear model. No SGBs reached significance at the MRM FDR-corrected p-values < 0.05 threshold.

**Fig. S3.**
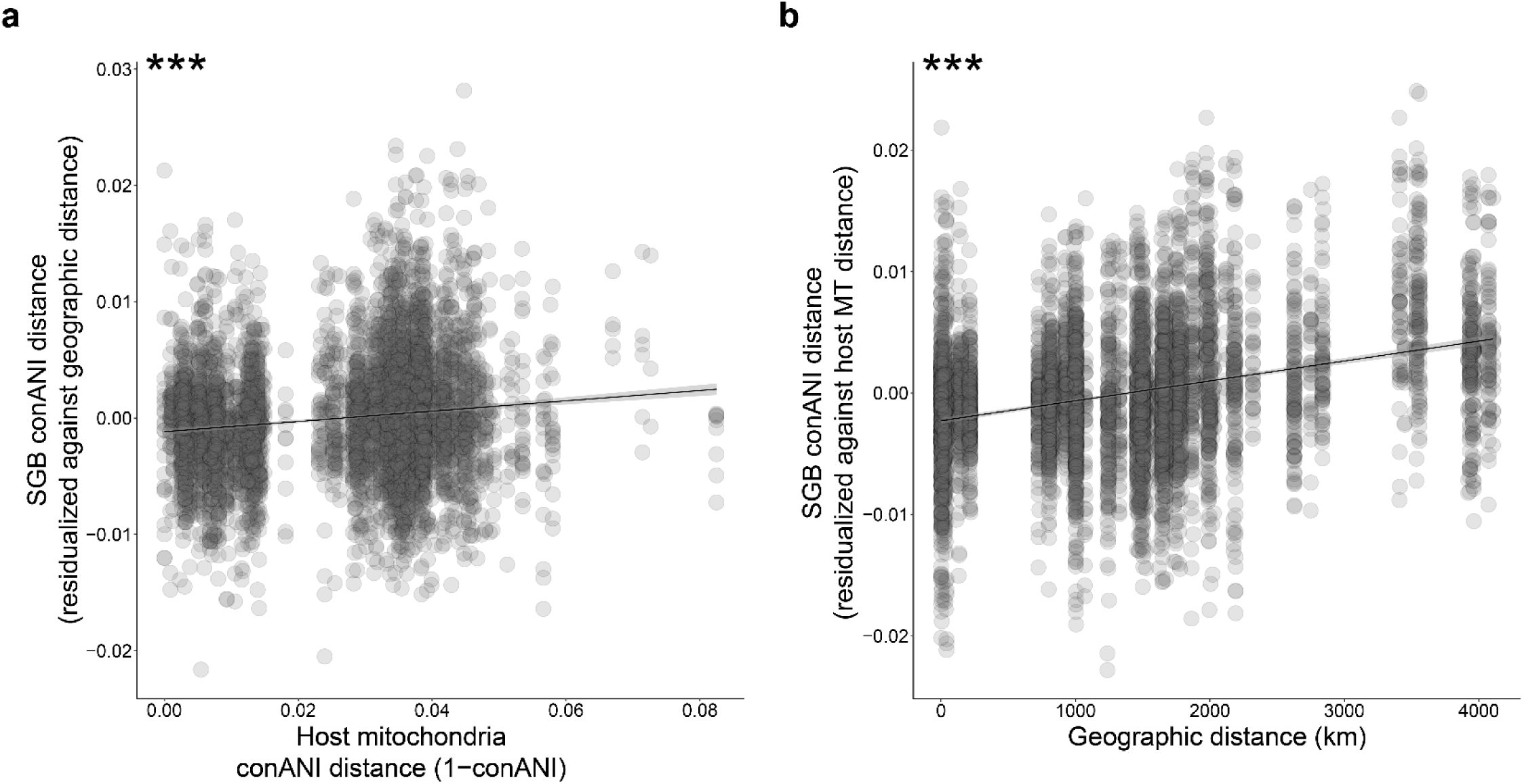
Aggregating gut bacterial strain comparisons across SGBs indicates widespread IBD and relatively weaker IBH. A) Scatter plot shows the relationship between strain genomic similarity within SGBs (1 – conANI) and geographic distance, after accounting for the effects of host MT genetic distance. Y-axis shows genomic similarity (1 – conANI) residualized against host MT genetic distances. B) Scatter plot shows the relationship between strain genomic similarity within SGBs (1 – conANI) and host MT genetic distance, after accounting for the effects of geographic distance. Y-axis shows genomic similarity (1 – conANI) residualized against geographic distances. In (A) and (B), trendlines and shading indicate best-fit linear regressions with standard errors. Asterisks indicate regression p-value; *** < 0.001.

**Fig. S4.**
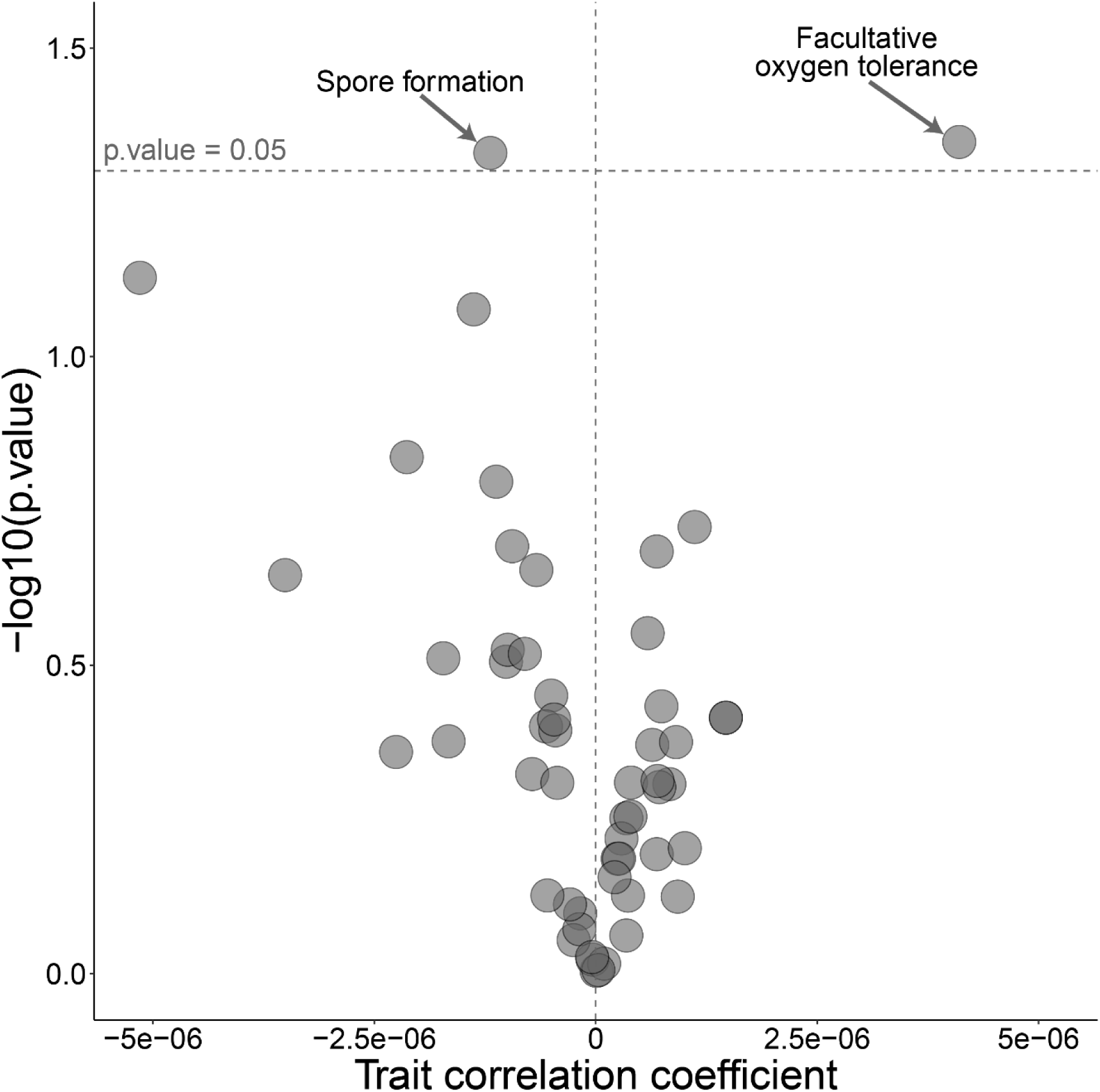
Traitar analyses confirm negative relationship between sporulation ability and strength of IBD. Volcano plot shows the significance (y-axis) of association between the presence of bacterial traits (points) and the strength of IBD. The strength of IBD was measured as the MRM Pearson correlation coefficient describing the relationship between geographic distance and strain genomic divergence between mice within SGBs residualized against host MT genetic distances.

